# Analysis of six-decadal seed mass and emergence records in mast species shows little inter-annual variability

**DOI:** 10.1101/2020.12.21.423701

**Authors:** Yang Liu, Yousry A. El-Kassaby

## Abstract

Patterns of crop production in mast species do not track crop-year climate, but instead are regulated by climate cues in prior-years. Whether the pattern of year-to-year seed mass variation is coupled in time with mast seeding, maintaining seed mass-number trade-offs, and coherently driven by similar climate cues as other seed traits (e.g. seed germination) remains unknown. Using ca. 6,000 long-term seed inventory data over the years 1955-2015 in conifers, this retrospective study revealed the temporal patterns of mast species’ seed mass and its associated trait, seed germination. To pinpoint their ecological drivers, pairwise correlation analysis was performed between each trait and seasonal climates in crop year and four prior-years. Using climate variables key to each trait, regression models were constructed to project trait values. Findings showed minor seed mass variation among years, which rejects the generality of seed mass-number trade-offs in many plant species. This result reasonably arises as the economies of scale (compensating benefits) theory are often used to account for mast seeding but not for seed mass. Moreover, final germination fraction also varied little over time, but exhibited an increasing tendency. In addition, we found that temperature-based climate variables drive seed mass, number, and germination variation, but these variables in different seasons of crop year or prior-years did not have equal influences on trait variability. Finally, regression models showed that the number of frost-free days and evapotranspiration are crucial to the three traits and climate in autumn is a critical season, followed by summer and winter. This study holds considerable promise for explaining reproductive strategies of taxonomic groups with mast seeding characteristics in allocating reproductive resources to different life-history traits using ecological signals.

## Introduction

Mast behaviour - synchronous and highly temporally variable production of fruits and seeds - is a crucial population process in plant species [1] with major cascading effects on ecosystems functioning [2-4]. Mast years - boom-bust seed production year - have been well documented in perennials, particularly in woody tree species such as many Fagaceae and Pinaceae (e.g. [5, 6]). A combination of climatic events is often used to explain inter-annual variation in crop production [7-10]. Numerous studies in tree species have demonstrated that mast years are associated with specific climate conditions before mast seeding [11, 12], and particularly with summer temperatures 1-2 years prior to mast [13-15].

Increased crop production (masting) prompts high fecundity (seed numbers). Within-year or lifetime trade-offs between seed mass and number have been documented but almost only in species that have no mast seeding (e.g. [16-18]). To our knowledge, only one example tested the seed mass-number trade-off in a masting species (oak) but only in a single year [19]. To date, no study has been conducted to investigate this relationship in mast species between mast and lean years. Whether such a negative relationship still holds in mast species over time is yet unknown. In general, trade-offs between fitness-enhancing traits arise when they are favoured by selection but compete for a limiting resource and/or share a genetic correlation [20] such as offspring size (or seed mass for plants) and number [18, 21]. For a given amount of resources available to a parent plant, seed mass and number, a priori, constrain one another and this constraint further affects individual plant fitness [22, 23]. Nonetheless, the economies of scale theory (e.g. predator satiation, increased pollination efficiency, and improved seed dispersal), pollen coupling hypothesis (i.e. endogenous resource dynamics associated with pollen limitation; reviewed in [2, 24]), and the environmental prediction hypothesis (i.e. favorable conditions for seedling establishment also conducive to mast seeding; [25, 26]) have been used to account for mast behaviour, but not for seed mass. In addition, seed mass variation often has concomitant effects on seed dormancy (i.e. delayed onset of seed emergence) [27] and both traits have been jointly investigated at ecological and/or evolutionary levels (e.g. [28-30]). Thus, seed mass, number, and dormancy (or germination) vary and likely evolve in a coordinated manner; however, their trade-offs are found not to be consistent among species (e.g. [31]). This implies different adaptive constraints on fitness for a given trait in different species [32].

This retrospective study used conifers as a study system to test whether mast species’ seed mass variability is commensurate with mast seeding variability and whether climate cues reported as key to masting (seed number) in previous studies also exert a significant effect on seed mass and germination variation. Based on a six-decadal (1955-2015) seed trait inventory data and climates in historical time series across British Columbia, Canada, we unveiled the trait variation patterns at temporal scales and performed correlation analysis between each trait and seasonal climates in seed crop year and four prior-years. Finally, we employed a machine-learning approach (stepwise model selection) to identify top important climate variables from those significantly correlated ones, whereby regression models were constructed to project trait values by using these key predictors.

## Material and Methods

### 1.1. Study system and data compilation

Three conifers were chosen, including white spruce (*Picea glauca*), Douglas-fir (*Pseudotsuga menziesii*), and lodgepole pine (*Pinus contorta*). “Spruce”, “Fir”, and “Pine”, if used, denote the three-study species, respectively. In general, conifers follow two types of reproductive cycles, for instance, spruce (*Picea* spp.) and Douglas-fir (*Pseudotsuga* spp.) have two- and most pine species (*Pinus* spp.) have three-year reproductive cycles. For white spruce, Douglas-fir, and lodgepole pine, heavy or good cone crops (masting) occur every *c*. 5-10, 3-12, and 1-3 years, respectively (see illustration in [33]).

In total, we used 1,314, 1,070, and 3,458 observations of seed mass and final germination fraction for white spruce, Douglas-fir, and lodgepole pine, respectively, over the period 1955-2015 across British Columbia, Canada. Sample depth in each of the three species was showcased in Fig. A.1-A. The seeds used for tests (e.g. seed mass, germination) had 97% purity and 4.9-9.9% moisture content on a fresh weight basis. Seed mass unit was gram per 1,000 seeds averaged over 3-5 replicates. Our data showed that the mean mass of Douglas-fir (10 g/1K seeds) is 3-5 times as heavy as that of white spruce and lodgepole pine (2.06 and 2.93 g/1K seeds, respectively). Final germination fraction averaged over three replicates was assayed after seed harvest.

Previous studies have demonstrated that the number of seed cones per tree (masting) is a positive correlate with seed number (e.g., [34, 35]), we therefore used seed number as a proxy for mast seeding. As we had no archived seed number data, we resorted to the literature and the correlations between seed number and climate were tested based on previous reports, including *Picea glauca* in Yukon, a territory in the north of British Columbia [36] and *Pinus ponderosa* in Boulder, Colorado [37]. To directly ascertain ecological drivers of seed number, we searched the literature and used total seed number per area in Engelmann spruce (a sister species with white spruce) from the southern Rocky Mountains [38] and seed number records (inter-annual masting variation) for the three conifers [39].

Climate data in historical time series were obtained in the seed crop year (year *i*) (records in our three focus species and Engelmann spruce from the literature) and four sequential prior-years (years *i*-1, *i*-2, *i*-3, and *i*-4) using ClimateNA ver.5.41 [40]. This software calculates more than 200 monthly, seasonal, and annual climate variables, such as maximum or minimum mean temperature [*T*_max_ or *T*_min_], evapotranspiration [*Eref*], number of frost-free days [*NFFD*], degree-days above 5°C or below 18°C [*DD* > 5 or *DD* < 18].

### 1.2. Data analysis

All data analyses were conducted in R ver.3.4.4 [41]. First, pairwise Pearson’s product-moment correlation analyses were performed between each of the three focal traits (i.e. seed mass, final germination fraction, and Engelmann spruce seed number from the literature) and seasonal climates in the temporal sequence (four-prior years until a seed crop, that is, years *i*-4 through *i*). All *p*-values for these comparisons were adjusted using the sequential Bonferroni correction [42]. We compared adjusted *p*-values to critical values calculated based on α = 0.05 so that fewer than 5% of variables identified as significant were false positives (or type I error), that is, *q*-values [43, 44]. Then, we tested for how well most important climate variables could be used for the trait projection. We fitted each of the three traits as a function of a set of key climate variables in a linear manner. The importance of each variable with significant correlations (both adjusted *p* and *q* < 0.05) was assessed by computing the difference between Akaike information criterion values (ΔAIC) of the models with or without the concerned variable – the higher a ΔAIC, the more the importance of the variable in the model. We completed the variable selection first by using a machine-learning approach in a stepwise algorithm (both forward and backward) in the “MASS” package in R [45] and then by choosing the top six relative important ones (to avoid overfitting in modelling) based on the *R*^2^ contribution averaged over orderings among regressors using the “relaimpo” package in R [46]. The predictor terms (all standardized) of the final models consisted of six most important climate variables and all their two-way interactions. The final models were validated by leave-one-out cross-validation (LOOCV).

## Results

The year-to-year seed mass variation and its 95% credible interval band were visualized in Figure 1A and their coefficients of variation (CVs) were ca. 11.3, 13.3 and 9.6% in Spruce, Fir, and Pine, respectively. This indicates that seed mass variation among years was small in the three conifers. Besides, seed mass variation was minor in space as well (no significant high density in each panel of Fig. A.1-B). In contrast, seed production CVs in mean crops among years were ca. 160, 150, and 90% in Spruce, Fir, and Pine, respectively (Fig. A.2). In addition, CVs in final germination fraction were not high among years (ca. 16.2, 14.7, and 8.8% for Spruce, Fir, and Pine, respectively; Fig. 1A) and showed an elevated tendency (Fig. 1A). This indicates that seed dormancy (germination-arrest mechanism) has tendency to decrease possibly due to climate change. Seed mass and final germination fraction had weak but significant correlations (Fig. 1B; for Spruce: *F*_1, 1312_ = 20.5, *p* < 0.0001; for Fir: *F*_1, 1089_ = 45.3, *p* < 0.0001; and for Pine: *F*_1, 3456_ = 251, *p* < 0.0001). Given that two key climate variables, July Δ*DD* > 5 (difference in July degree-days above 5°C in two prior-years) and July Δ*T*_max_ (July maximum temperature difference in two prior-years), are positively correlated with white spruce masting [36], both variables were negatively correlated with seed mass (Fig. 1C-a; for July Δ*DD* > 5 and mass relationship: *F*_1, 1312_ = 24.5, *p* < 0.0001; for July Δ*T*_max_ and mass relationship: *F*_1, 1312_ = 32.6, *p* = 1.4e-08). Given that spring *T*_ave_ (spring mean temperature 2 years before seed crop) is negatively correlated with pine seed masting [37], it was positively correlated with seed mass (Fig. 1C-b; *F*_1, 3450_ = 155, *p* < 0.0001). This indicates that climate variables significantly correlated with seed number variability also affect seed mass but in an opposite manner in conifers. However, the effect of these key climate variables for masting was minor on seed mass, as shown by a 10-degree change in July Δ*DD* > 5 leading to less than a 1-g change in 1K-seed mass (Fig. 1C). In contrast, climatic effects on seed number are substantial with a 1-degree temperature difference between two preceding summers before seed crop generating a 10-fold change in seed number [14].

**Figure 1.**
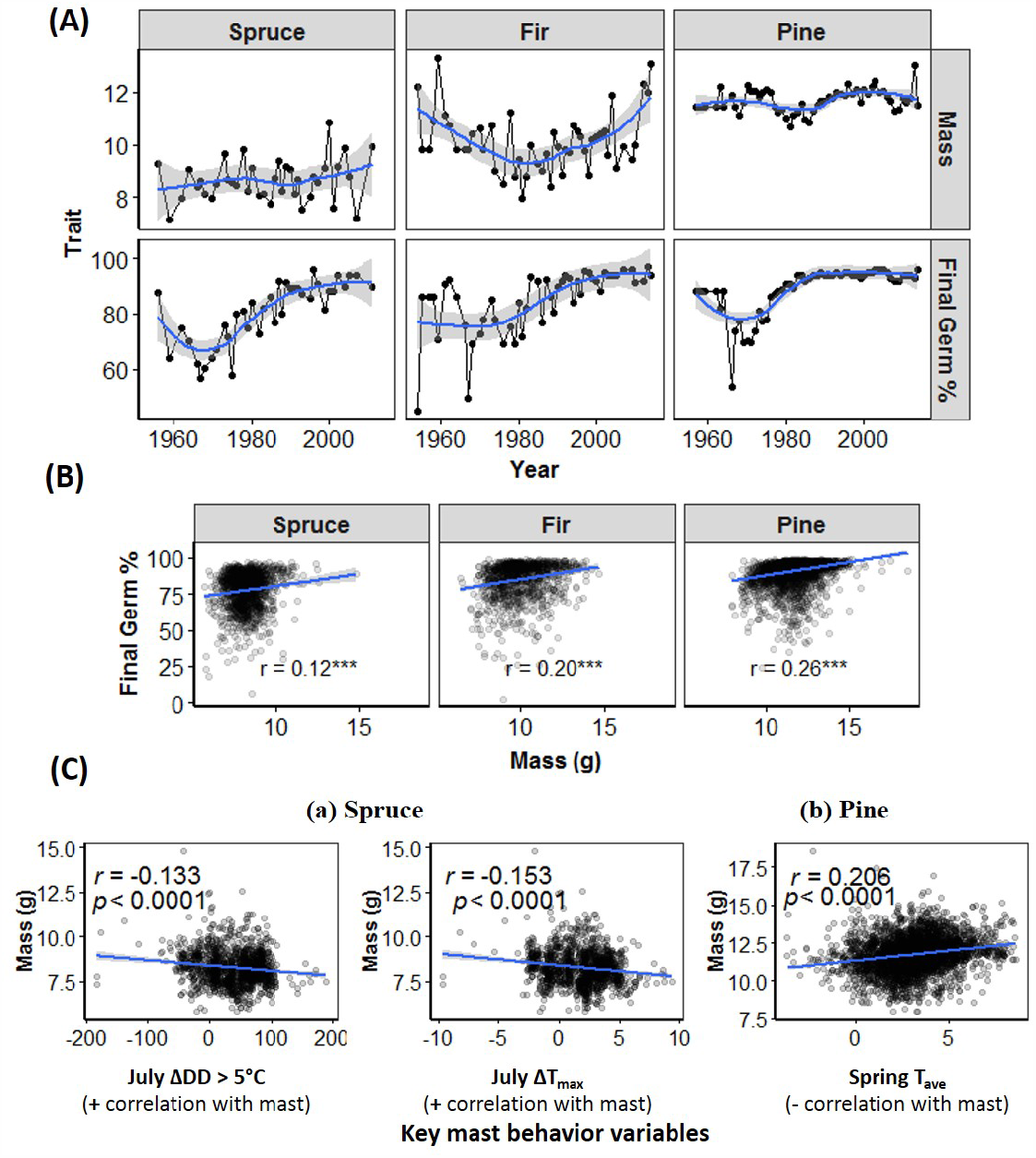
Changes in seed mass and final germination fraction for three conifer species spanning the years 1955-2015 (A); Pearson’s correlation *r* between seed mass and germination fraction by species (B) and between seed mass and climatic variables of mast seeding (C).

Based on the literature, important conifer seed-set periods are related to pollen and seed development. The top climatic drivers for the three traits were mainly temperature-based (showcased in Fig. 2 y-axis). Of these environmental drivers, only *DD* < 18 had an inverse correlation with both conifer seed mass and germination (Fig. 2A and 2B). Moreover, different time periods were of disparate importance to seed mass between species. Fir seed mass was highly correlated with climates throughout crop year and 4 prior-years, especially highlighting summer and autumn seasons (Fig. 2A). Summer climate in crop year and three prior-years was important to Pine seed mass (Fig. 2A). Spruce seed mass had a relatively low relatedness with climate (Fig. 2A). By contrast, spring climates were more correlated with final germination fraction than autumn counterparts in Spruce and Fir (Fig. 2B); Pine seed germination was significantly correlated with climates of all four seasons throughout years *i* to *i* −4 (Fig. 2B).

**Figure 2.**
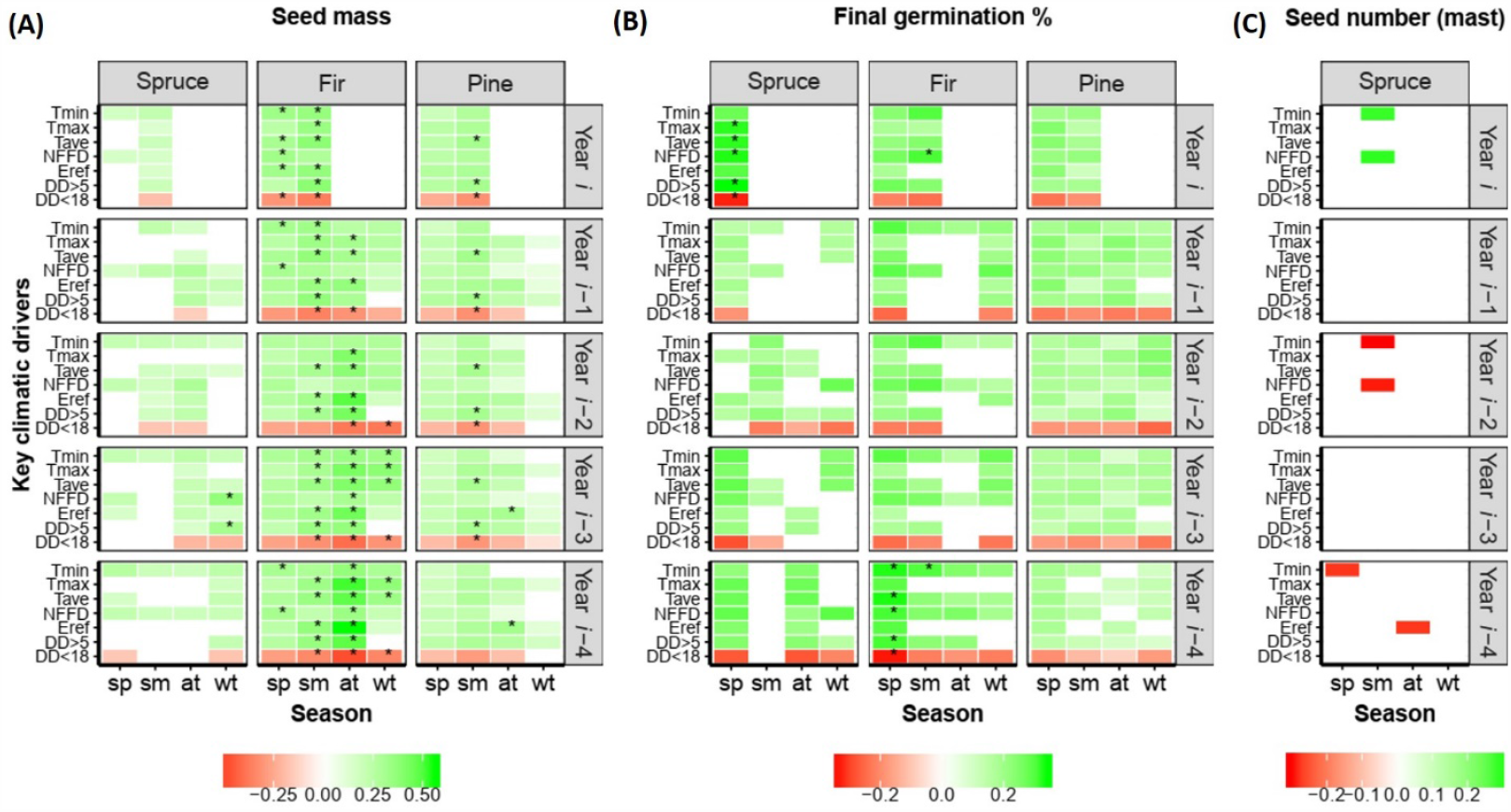
Pearson’s correlation *r* between traits and seasonal climatic variables (A-C). Abbreviations for climate variables: minimum mean temperatures (*T*_min_), maximum mean temperatures (*T*_max_), mean temperature (*T*_ave_), the number of frost-free days (*NFFD*), evapotranspiration (*Eref*), degree-days above 5°C (*DD* > 5), and below 18°C (*DD* < 18). *: correlations |*r*| ≥ 0.3 (∼ top 3%) with significance at *p* < 0.0001 (sequential Bonferroni-corrected) and *q* < 0.0001. Non-significant correlations (adjusted *p* or *q* ≥ 0.05) are not displayed (blank). Given seed harvest in the autumn of year *i*, no correlation analysis was performed (blank).

Each black dot in (A) is a median trait value in time series (spruce and pine seed masses magnified by 4 times for visualization convenience) and blue trend line curves are smoothed with grey shading for 95% confidence interval bands using a loess method. The unit for seed mass and germination is g/1K seeds and %, respectively. Note that most updated knowledge indicate that the key climatic variables for *Picea* masting include difference in July degree-days above 5°C in two prior-years *i*-1 and *i*-2 (July Δ*DD >* 5°C) and difference in July maximum temperature in two prior-years (July Δ*T*_max_). In *Pinus*, the key climatic variable is spring mean temperature 2 years before seed cone maturation (Spring *T*_ave_). As of September 2020, there was no masting study conducted in *Pseudotsuga* spp.

Prior research has shown that critical seed-set periods affect mast seeding, including pollen and ovule meiosis [37], bud primordia development [13, 25], and pollination and fertilization [10, 38]. We found that *T*_min_ and *NFFD* in the summer of years *i* and *i* −2 were significantly correlated with spruce seed number (Fig. 2C). However, these two climate variables had opposite correlations with seed number in year *i* compared with year *i* −2 (Fig. 2C). Similar studies supported that summer climate in year *i* −2 was the most significant predictor of Spruce cone crops [36] and spring *T*_ave_ in year *i* −2 was the best predictor of Pine cone production [37].

Furthermore, based on top six climate variables key to each of the three traits, we constructed regression models to explain the variability of each trait. All the models performed fairly well and model residuals generally showed no systematic bias and patterns (Fig. A.3). While the LOOCV was successful for Fir seed mass (*r* = 0.58) and spruce seed number (*r* = 0.67) (Fig. 3), consistent with their high correlations with climate (Fig. 2A-C), LOOCV was less successful for the other traits (average *r* = 0.377; Fig. 3). In the final models, 42 climate predictors (i.e. six climate variables per model × seven final models) were chosen, where 28.6% of the predictors were *NFFD*, followed by *Eref* (19%). Moreover, of these predictors, 13 were summer-based (31%), followed by autumn and winter (∼ 25% each). Interestingly, except one-prior year (*i* −1), all the other years (i.e. crop year *i* and prior-years *i* −2 through *i* −4) were equally highly used in the models (∼23% for each of the four years). This indicates that one year preceding a crop year is likely less important to the variability of these traits than crop year and 2-4 prior-years in conifers, contrasting with previous findings in other mast species.

**Figure 3.**
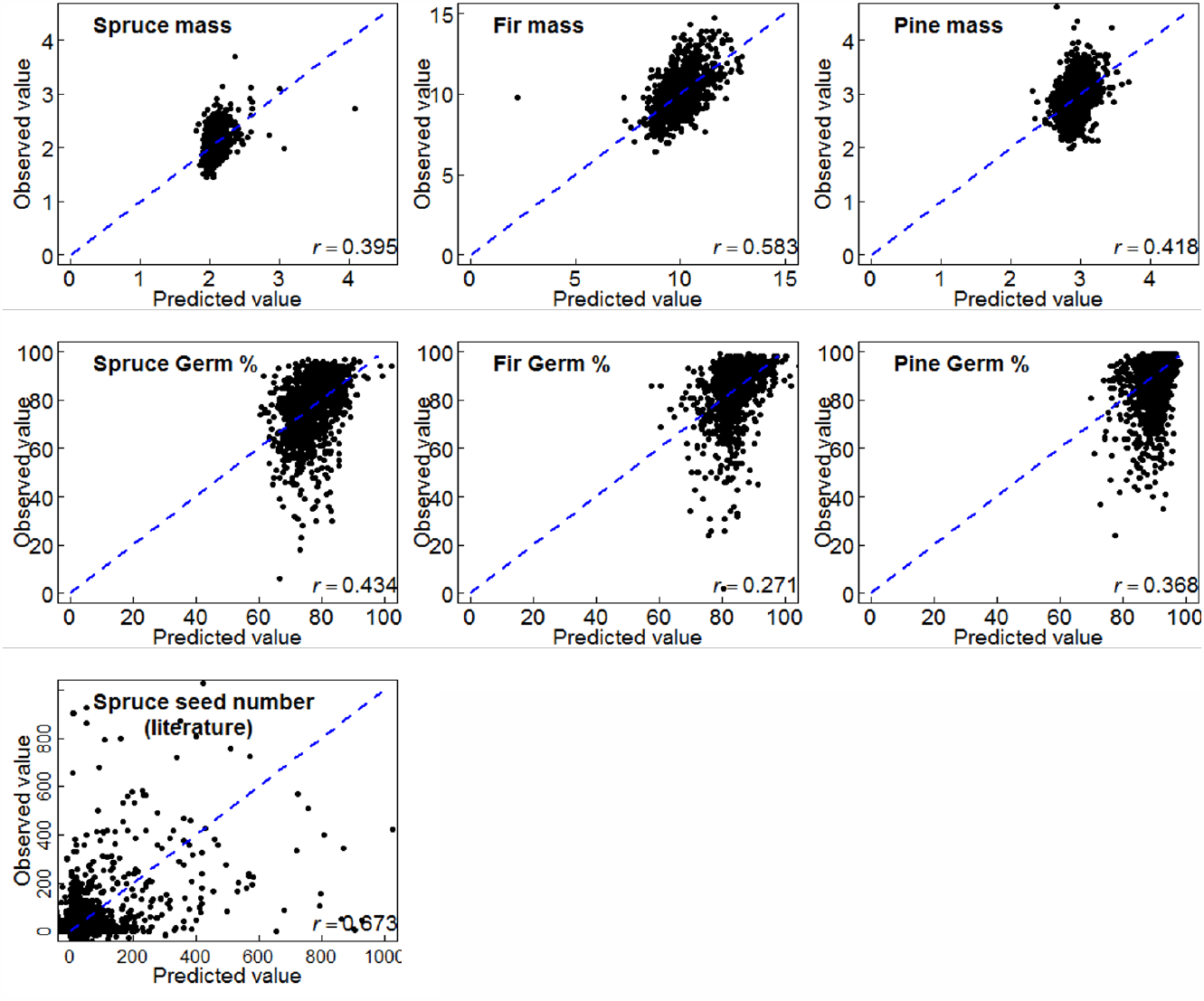
Leave-one-out cross-validation for trait projection models The dashed diagonal line represents the perfect match between predicted and observed values.

## Discussion

Mast species irregularly store energy for one or multiple years and the energy is then used for reproduction, such as flowering and fruiting [6, 47]. As such, mast seeding is the consequence of resource storage and allocation to reproduction, in combination with synchronizing mechanisms [48, 49]. Given that plants must accumulate a threshold amount of resources to initiate reproduction and this threshold cannot be attained in one season, plants adopt a strategy to accumulate resources over multiple years, which then translates into intermittent crop production by individual plants [48]. This formalizes one of the hypotheses to explain why many perennial plant species show intermittent reproductive cycles, for instance, large annual variation in seed number results in the production of large seed crops during some years, interspersed with other years of low seed crops [1, 5, 50, 51].

Further, this study showed minor year-to-year variation in seed mass and germination, indicating that seed mass and germination have no reproductive trade-offs with mast seeding. The same key seasonal climatic drivers had different importance to different species (different correlations shown in Fig. 2A), indicating that reproductive strategies are phylogenetically-dependent. This may be systematically associated with the mechanisms underpinning adaptive processes, ecological filtering, niche breadth, and internal cycling (e.g. turnover or generation time, reproductive cycle duration, inherent reproductive capacity, etc.) Through photosynthesis, assimilated carbon into non-structural carbohydrates (source activity) is partitioned to several sinks, including respiration, growth, reproduction, storage, and defence [52]. Moreover, trees in mass-flowering years are in a particular physiological status (e.g. low starch and elevated hormone levels; [53, 54]), which acts to prevent the floral induction process [55]. As such, changes in ecophysiological conditions during masting vs. non-masting seed-set periods may contribute to the variation in seed characteristics. Nonetheless, it is worth noting that many studies point to little relatedness between starch status (or non-structural carbohydrates) and masting seeding events (e.g. [56]). Endogenous plant cycles (e.g. via hormone fluxes) considerably affect cone-bud initiation and gender differentiation in conifers [57], which involves interactive roles of the environment, including photoperiod durations [57]. This suggests whether the three-study species have distinctive hormone regulations may be a logical entry to further study species-specific disparities.

Finally, it is interesting to consider crown dynamics in the mast behaviour. From a lifetime perspective, aging has negative effects on the fructification and net cone production decreases with crown dynamics over time (see [58] for crown dynamics patterns). As crown lifts, crown size decreases and thus cone-bearing surface decreases. However, at what age such canopy attribute related with cone-bearing begins to hold needs further investigations.

## Conclusion

There is no clear seed mass-number trade-off in coniferous mast species. Final seed germination fraction shows little variation over time, but has an increasing tendency possibly as the consequence of climate change. Regression models indicate that the number of frost-free days and evapotranspiration are key to seed mass, number, and germination; autumn is a critical season for shaping these traits, followed by summer and winter. Overall, this study reveals what and how ecological signals drive the allocation of resources to life-history traits at the plant-to-seed transition.

## Acknowledgements

We are grateful to the staff affiliated to British Columbia Tree Seed Centre (TSC; Surrey, Canada) for providing their long-term historical records and to the scientists who contributed to the seed number data on which this analysis is based. This project was funded by the Johnson’s Family Forest Biotechnology Endowment and the National Science and Engineering Research Council of Canada Discovery and Industrial Research Chair to Y.A.E.

## Appendix A

**Figure A.1.**
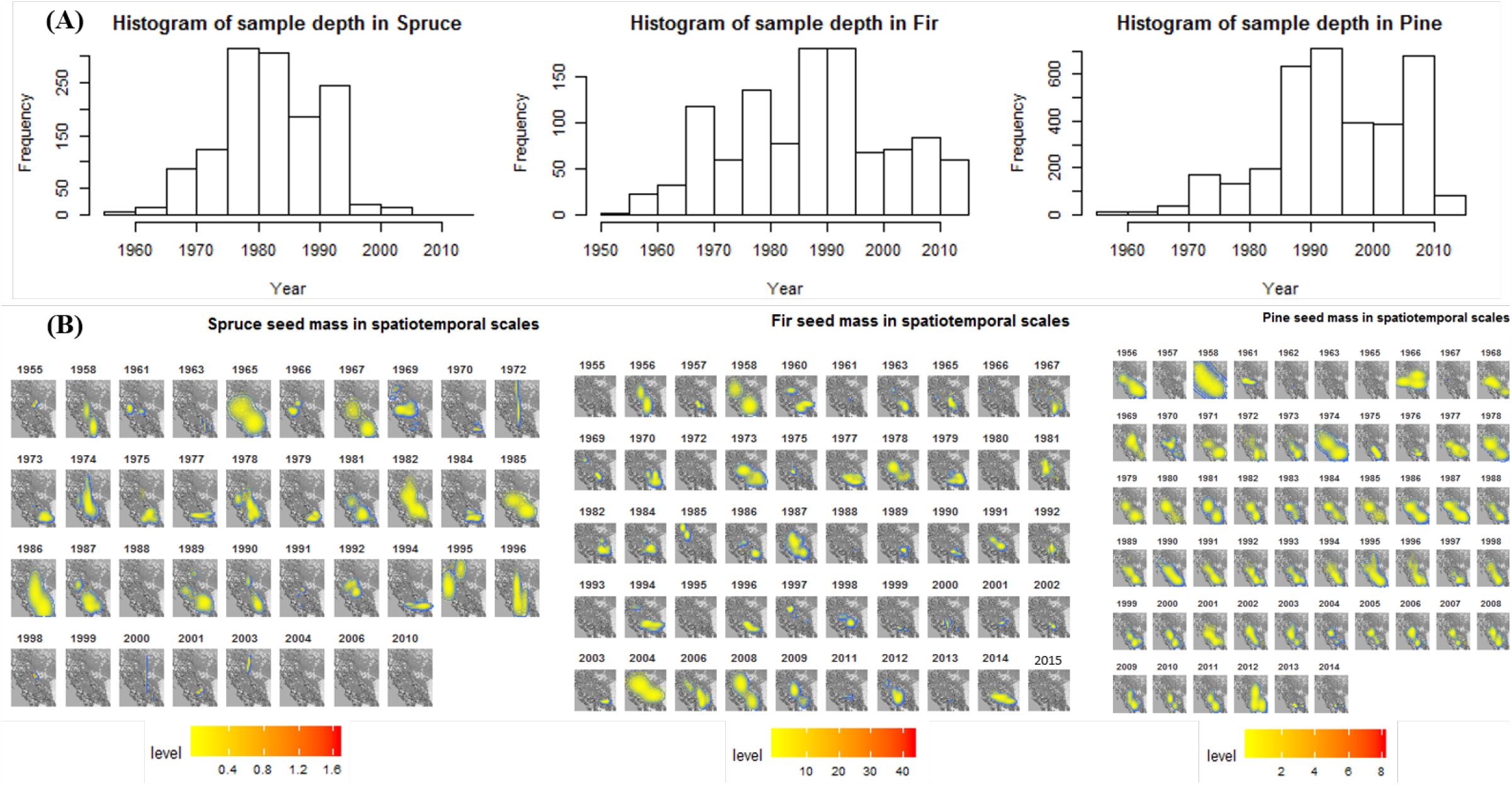
Sample depth (A) and seed mass contour and density 2-D plot (B) for the three-conifer species by year Level bar in (B) indicates the density of seed mass within geospatial areas displayed.

**Figure A.2.**
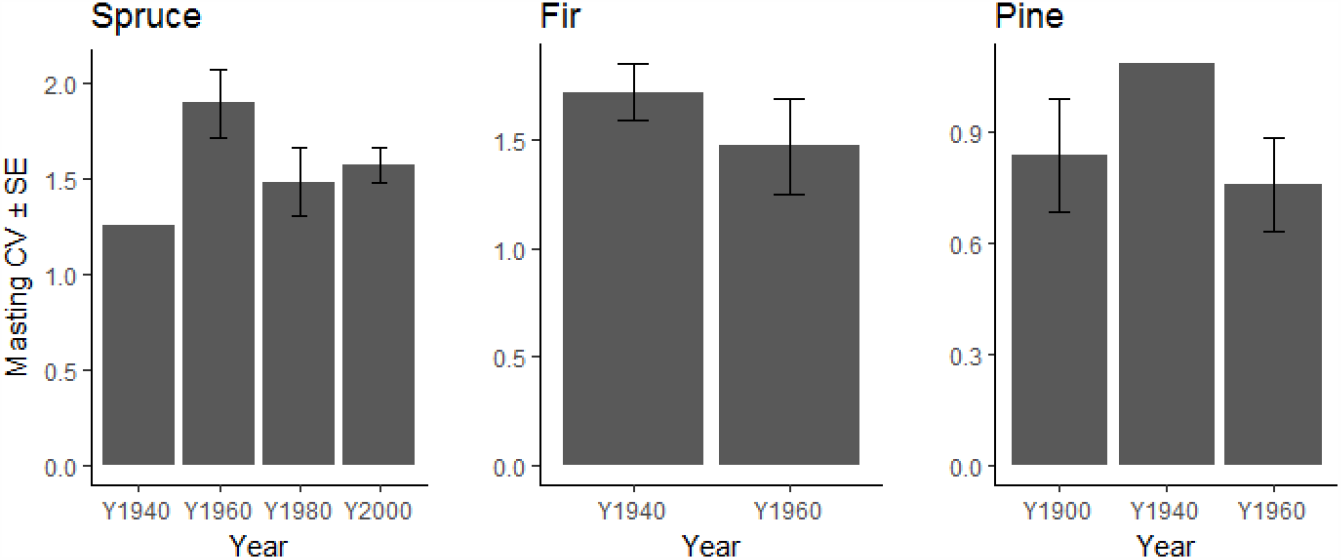
Seed masting records in 20-year time periods Plants have continuous variation in the population-level coefficient of variation (CV) and standard errors (SE) were calculated based on the records from different studies. All pairwise Wilcoxon tests for masting CV between time periods were not significant at α = 0.05, indicating no decadal trend.

**Figure A.3.**
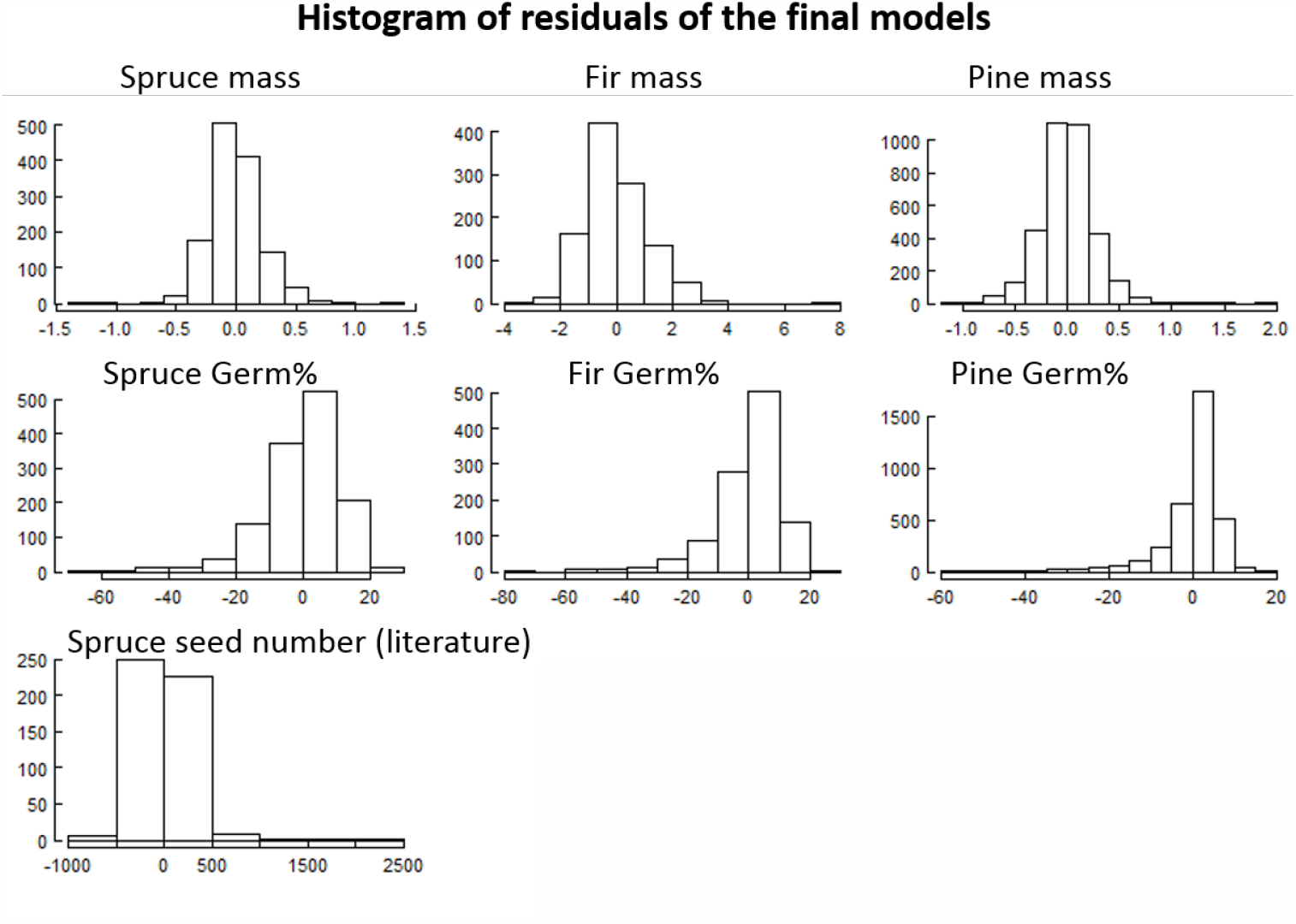
Residuals of all final models for seed mass, germination, and number in three focal species Each model includes six selected climate variables and all their two-way interactions.

